# NR4A3 deficiency ameliorates contact hypersensitivity and exacerbates psoriasis by regulating gene expression in dendritic cells

**DOI:** 10.1101/2024.02.20.581116

**Authors:** Mayuka Katagiri, Naoto Ito, Kazuki Nagata, Shiori Nakano, Natsuki Minamikawa, Niya Yamashita, Miki Takahashi, Masami Kurihara, Masanori Nagaoka, Miki Ando, Tomoka Ito, Takuya Yashiro, Akihiko Yoshimura, Chiharu Nishiyama

## Abstract

NR4A3 is a transcription factor that belongs to the nuclear receptor superfamily. To reveal the roles of NR4A3 in skin diseases, we constructed contact hypersensitivity (CHS)- and imiquimod-induced psoriasis models in *Nr4a3*^-/-^ mice. In the CHS induced *Nr4a3*^-/-^ mice, ear swelling was significantly reduced, accompanied by suppressed migration of dendritic cells (DCs), in which CCR7 expression was reduced. In contrast, ear swelling in psoriasis model *Nr4a3*^-/-^ mice was enhanced. The expression levels of inflammatory cytokines in psoriasis skin lesions and in TLR7-ligand-stimulated DCs were increased by NR4A3 deficiency. Subcutaneous (s.c.) injection of hapten-treated *Nr4a3*^+/+^ DCs induced ear swelling in *Nr4a3*^-/-^ mice, and s.c. injection of TLR7-ligand-stimulated *Nr4a3*^-/-^ DCs caused significant ear swelling. Compared with *Nr4a3*^+/+^ DCs, *Nr4a3*^-/-^ DCs expressed lower levels of CCR7 and IRF4, and higher levels of TLR7, IRF7, IFN-β, and PU.1. Taken together, NR4A3 deficiency ameliorates CHS and exacerbates psoriasis by regulating the expression of CCR7, TLR7, and related transcription factors in DCs.

## Introduction

NR4A3/NOR-1 belongs to the nuclear receptor (NR) superfamily. Although almost all NR members, including steroid hormones, fat-soluble vitamin receptors, and PPARs, are activated by the binding of their ligand molecules to a pocket in the ligand-binding domain (LBD), NR4A3 and other NR4A subfamily members (NR4A1/NUR11 and NR4A2/NURR1) are orphan receptors. A 3D-structure study of NR4A2 indicated that NR4A subfamily members are constitutively activated and regulate gene expression in a ligand-independent manner because the pockets of their LBDs are too small for ligand binding^1^.

The role of NR4A3, also termed neuron-derived orphan receptor 1 (NOR1), was first described in neurons, and *Nr4a3* gene-disrupted mice exhibited bidirectional circling behavior due to defects in the development of the semicircular canals of the inner ear^2,^, and in postnatal development of the hippocampus^2,3^.

In addition, the involvement of NR4A3 in circulatory system disease has been reported, in brief, high expression of NR4A members was observed in human neointimal smooth muscle cells (SMCs) of atherosclerotic lesions, and protective roles of NR4A members in atherosclerosis were indicated in a study using transgenic mice expressing dominant-negative NR4A1 or full-length NR4A1 in SMCs^4^. In line with this study, an investigation utilizing *Nr4a3*^-/-^ *Apoe*^-/-^ mice revealed that NR4A3 deficiency reduced atherosclerosis^5^. In contrast, a bone marrow transplantation experiment in which irradiated *Apoe*^-/-^ mice were reconstituted with NR4A3-deficient hematopoietic stem cells showed that NR4A3 deletion in hematopoietic stem cells accelerated high-fat diet-induced atherosclerosis formation and macrophage recruitment^6^, suggesting a hematopoietic cell-specific function of NR4A3.

In terms of hematopoietic lineages, a great deal of research has been conducted on the combined functions of all NR4A family members in T cells. T-cell-specific triple knockout of NR4A receptors induced lethal systemic autoimmunity in mice, along with a deficiency in Treg development^7^. A study utilizing the transfer of CAR-T cells against human B-cell malignancies into human tumor-bearing mice demonstrated that CAR-T cells lacking all three NR4A transcription factors, which are characteristic of CD8^+^ effector T cells and do not enter the exhausted state, promoted tumor regression^8^. In contrast to the complementary functions of the NR4A family in T cells, the single suppression of NR4A3 had several impacts on monocyte lineages. In human alternative macrophages, silencing NR4A3 decreased the expression of M2 markers^9^. In murine dendritic cells (DCs), in which NR4A3 expression is increased during stem cell development and upon TLR-mediated stimulation, knockdown of NR4A3 inhibited the TLR-mediated activation of DCs ^10^. A deficiency in NR4A3 reduces the migration of DCs and DC-mediated immune responses that are activated by infectious gram-negative bacteria^11,12^.

DCs play key roles in innate and adaptive immune responses by providing a first line of defense against infection. In the present study, we constructed two kinds of DC-related skin inflammatory disease models, contact hypersensitivity (CHS) and psoriasis, in NR4A3 knockout (KO) mice, to clarify the roles of NR4A3 in skin homeostasis and the function of DCs in the skin.

## Results

### Effects of NR4A3 deficiency on CHS

To evaluate the roles of NR4A3 in skin diseases, we first constructed a CHS model in *Nr4a3* KO mice. As shown in **Fig. 1a**, the degree of ear swelling after FITC challenge in FITC-sensitized *Nr4a3^-/-^* mice was significantly lower than that in *Nr4a3^+/+^* mice. Suppression of ear swelling in *Nr4a3^-/-^* mice was also observed in CHS caused by the administration of low-dose DNFB as a hapten (**Supplementary Fig. S1a**). We analyzed DCs in the cLN by flow cytometry and found that the frequency and absolute number of migDCs were markedly decreased in *Nr4a3^-/-^* mice treated with FITC (**Fig. 1b**) and other haptens (**Supplementary Fig. S1B**), whereas resDCs were either moderately reduced (**Fig. 1b**) or unaffected (**Supplementary Fig. S1b**). Moreover, the FITC-induced increase in the number of memory CD4^+^ T cells and effector CD8^+^ T cells in the cLN was significantly reduced in *Nr4a3^-/-^* mice (**Fig. 1c**). The hapten-induced increase in effector CD8^+^ T cells in the cLN of *Nr4a3^-/-^* mice was also suppressed in other CHS models (Supplementary **Fig. S1c**).

**Figure 1.**
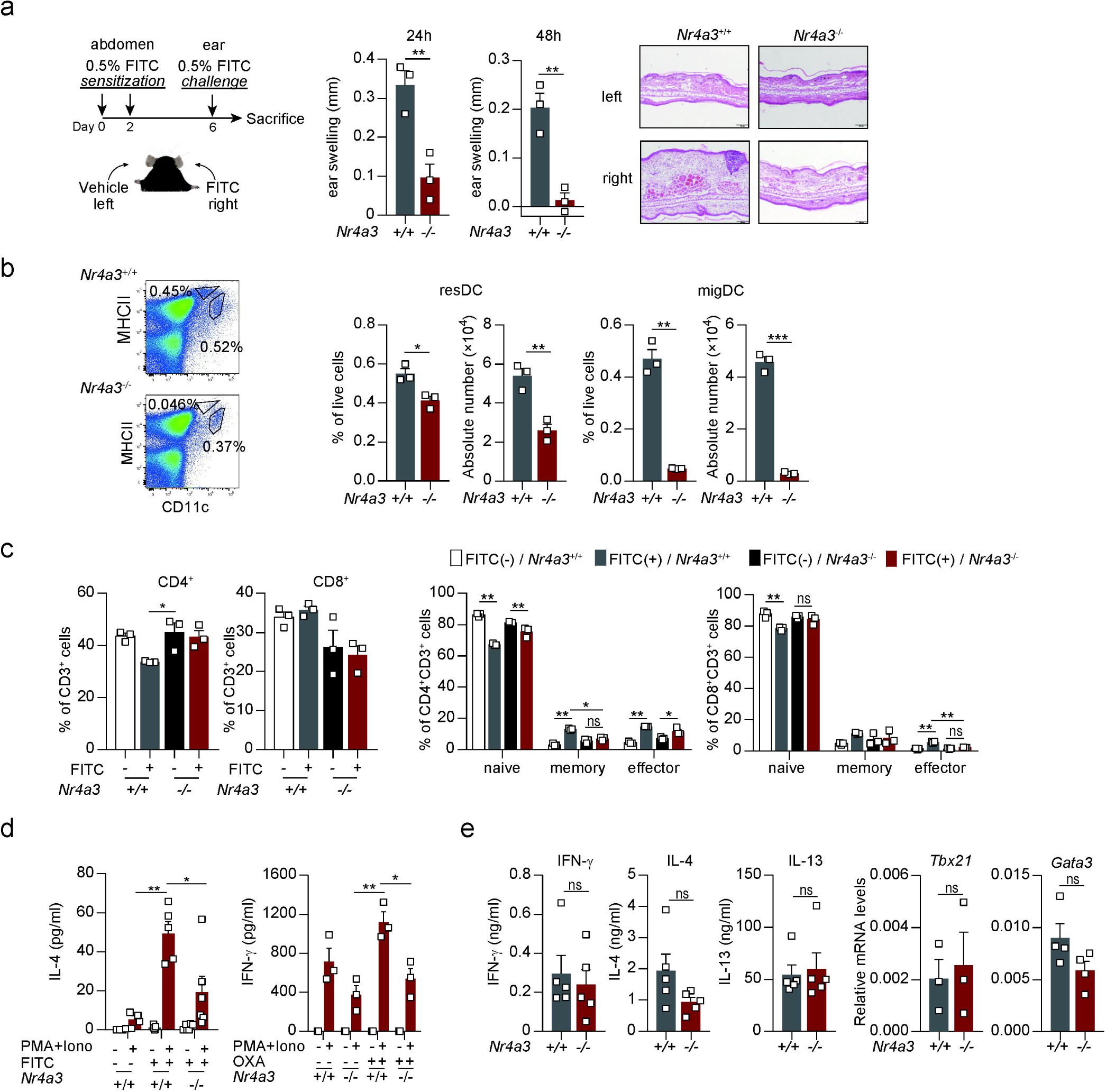
CHS is ameliorated in *Nr4a3^-/-^* mice. **a.** Schematic of the schedule used to establish the FITC-induced CHS model (left). Ten-week-old male *Nr4a3^+/+-^* and *Nr4a3^-/-^* mice were sensitized by abdominal application of 400 μL of 0.5% FITC on Days 0 and 2, and challenged by application of 20 μL of 0.5% FITC (right ear) or vehicle (left ear). Ear thickness in CHS-induced mice was measured 24 and 48 h after challenge (center). (Ear swelling) = (ear thickness 24 and 48 h after the challenge) – (ear thickness just before the challenge). H&E staining profiles of the ear (right). **b.** The proportion and absolute number of resDCs and migDCs in the cLNs harvested 48 h after FITC challenge were determined via flow cytometry with the gating strategy shown in **Supplementary Fig. S1b**. **c.** Proportions of CD4^+^ and CD8^+^ T cells in the cLN 48 h after FITC challenge determined via flow cytometry using the gating strategy shown in **Supplementary Fig. S1c**. **d.** Left: The amount of IL-4 produced by CD4^+^ T cells from FITC-sensitized or nonsensitized mice. Mice were sensitized with FITC on Days 0 and 2 and sacrificed on Day 6 without challenge. CD4^+^ T cells isolated from the cLNs were incubated in the presence or absence of 50 ng/mL PMA and 1 μg/mL ionomycin for 24 h, after which the culture supernatants were collected to determine the concentration of IL-4. Right: The amount of IFN-γ produced by CD4^+^ T cells from OXA-sensitized mice. Sensitization was performed according to the schedule shown in **Supplementary Fig. S1a**, and stimulation of CD4^+^ T cells was performed as described for the experiment shown in Fig. 1d (**left**). **e.** In vitro development of Th1 and Th2 cells. Naïve CD4^+^ T cells isolated from the spleen were incubated in plates coated-with anti-CD3 and anti-CD28 Abs under the Th1- or Th2-polarizing conditions for 3 days. After additional incubation for 24 h in the presence of 50 ng/mL PMA and 1 μg/mL ionomycin, the concentrations of IFN-γ (Th1), IL-4 and IL-13 (Th2) in the culture media were determined via ELISA. Messenger RNA levels of *Tbx21* (Th1) and *Gata3* (Th2) in CD4^+^ T cells after 3 days of culture under CD3/CD28-stimulated and Th1/2-polarized conditions. All of the results in Fig. 1 are shown as the mean ± SE. Significance was determined by two-tailed unpaired Student’s *t*-test (**a**, **b**, **e**), or one-way ANOVA with the Tukey-Kramer multiple comparison test (**c**, **d**). \**p*<0.05; \*\**p*<0.01; \*\*\**p*<0.001; \*\*\*\**p*<0.0001; ns, not significant.

When CD4^+^ T cells isolated from the cLNs of the *Nr4a3*^+/+^ mice in the sensitized phase were stimulated with phorbol ester and ionomycin, an increased amount of IL-4 (FITC) or IFN-γ (OXA) was released into the culture supernatant, whereas cytokine production by CD4^+^ T cells was significantly reduced in *Nr4a3*^-/-^ mice (**Fig. 1d**). In contrast, the development of Th1 and Th2 cells from naïve CD4^+^ T cells under polarizing conditions was not affected by NR4A3 deficiency (**Fig. 1e**).

### Effects of NR4A3 on the function of DCs in CHS

We detected FITC-bearing DCs in the cLN of FITC-sensitized mice to demonstrate the effects of NR4A3 deficiency on the migratory activity of DCs (**Fig. 2a**). As shown in the dot-plot profiles, more than 3% of the FITC-positive cells were detected in the cLNs of *Nr4a3^+/+^* mice at 24 hours after FITC application; these cells were identified as DCs by the expression of MHCII^+^/CD11c^+^. In contrast, under the same conditions, FITC-positive cells were rarely detected in the cLNs of *Nr4a3^-/-^* mice, indicating a defect of in the migratory activity of *Nr4a3^-/-^* DCs.

**Figure 2.**
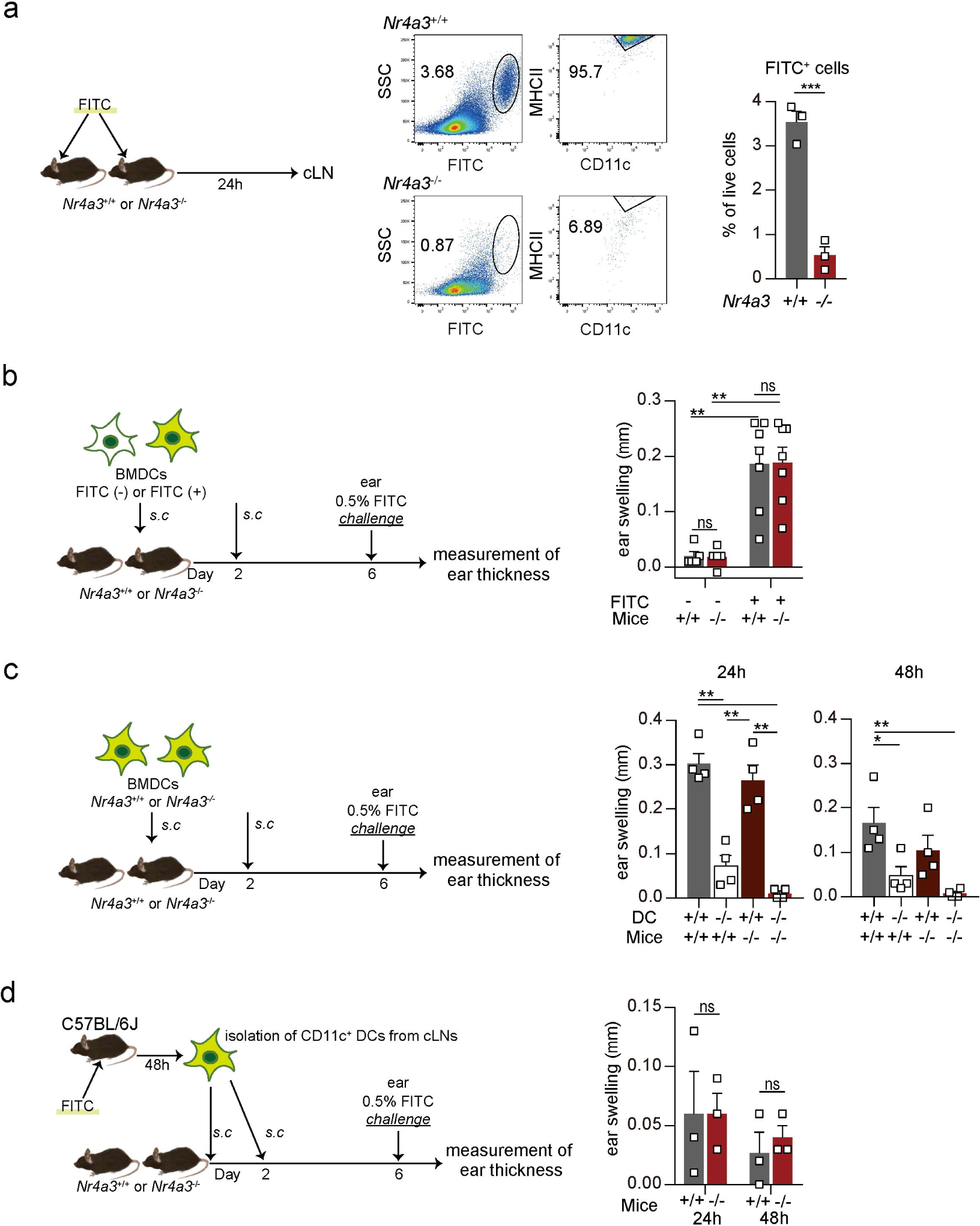
Effects of NR4A3 on the function of DCs in CHS. **a.** Migration of FITC-bearing DCs into the cLNs. Left: Schematic of the experimental procedure. The necks of 11-week-old male *Nr4a3^+/+^* and *Nr4a3^-/-^* mice were treated with 20 μL of 0.5% FITC, and the whole cells in the cLNs collected 24 h after FITC application was analyzed via flow cytometry. Right: The proportion of FITC-positive cells among the total live cells isolated from the cLNs is shown. **b.** Transfer of FITC-treated wild-type BMDCs into *Nr4a3^+/+^* and *Nr4a3^-/-^* mice. Left: Schematic of the experimental procedure. C57BL/6 BMDCs were incubated in the presence or absence of 0.025% FITC in culture media for 30 min. After the application of 400 μL vehicle to the neck skin of *Nr4a3^+/+^* or *Nr4a3^-/-^* mice, FITC-treated or nontreated BMDCs (5 × 10^5^ cells) were s.c. injected via the neck on Days 0 and 2. All of the mice were treated with 20 μL of 0.5% FITC in the right ear on Day 6, and the thickness of the ears was measured on Day 7. Right: Ear swelling of the right ear on Day 6 is shown. FITC (-); mice injected with FITC-free BMDCs, FITC (+); mice injected with FITC-treated BMDCs, +/+; *Nr4a3^+/+^*mice, -/-; *Nr4a3^-/-^* mice. **c.** Transfer of FITC-treated *Nr4a3^+/+^* and *Nr4a3^-/-^*BMDCs into *Nr4a3^+/+^* and *Nr4a3^-/-^* mice. Left: Schematic of the experimental procedure. *Nr4a3^+/+^* or *Nr4a3^-/-^*BMDCs incubated with 0.025% FITC for 30 min were s.c. injected into vehicle-treated *Nr4a3^+/+^* or *Nr4a3^-/-^* mice via the neck on Days 0 and 2. All mice were challenged with 20 μL of 0.5% FITC in the right ear and 20 μL of vehicle in the left ear on Day 6, and the thickness of the ears was measured 24 h and 48 h after challenge. Right: Ear swelling of the ears. DC +/+; FITC-treated *Nr4a3^+/+^* BMDCs were transferred to mice, DC -/-; FITC-treated *Nr4a3^-/-^* BMDCs were transferred to mice, Mice +/+; *Nr4a3^+/+^* mice were used as recipients, Mice -/-; *Nr4a3^-/-^* mice were used as recipients. **d.** Transfer of CD11c^+^ DCs isolated from the cLNs of FITC-treated wild-type mice into *Nr4a3^+/+^* and *Nr4a3^-/-^* mice. Left: Schematic of the experimental procedure. Four hundred microliters of 0.5% FITC was applied on the abdominal skin of C57BL/6 mice, and the mice were sacrificed to isolate CD11c^+^ DCs from the cLNs 48 h after application. The neck skin of *Nr4a3^+/+^* or *Nr4a3^-/-^* mice was treated with 400 μL of vehicle and s.c. injected with CD11c^+^ DCs (1 x 10^5^ cells) from FITC-treated mice on Days 0 and 2. All of the mice were challenged with 20 μL of 0.5% FITC in the right ear on Day 6, and the thickness of the ears was measured 24 h and 48 h after challenge. Mice +/+; *Nr4a3^+/+^* mice were used as recipients, Mice -/-; *Nr4a3^-/-^* mice were used as recipients. All of the results in Fig. 2 are shown as the mean ± SE. Significance was determined by two-tailed unpaired Student’s *t*-test (**a**, **b**, **d**), or one-way ANOVA with the Tukey-Kramer multiple comparison test (**c**). \**p*<0.05; \*\**p*<0.01; \*\*\**p*<0.001; \*\*\*\**p*<0.0001; ns, not significant.

We s.c. injected *Nr4a3^+/+^* BMDCs, which were incubated in the presence or absence of FITC, into the necks of *Nr4a3^+/+^* or *Nr4a3^-/-^* mice, and after the application of FITC to the ears of the BMDC-injected mice, we measured the thickness of the ears (**Fig. 2b left**). As shown in **Fig. 2b** (**right**), preinjection of *Nr4a3^+/+^* BMDCs carrying FITC into *Nr4a3^-/-^* mice induced obvious ear swelling comparable to that in *Nr4a3^+/+^* mice after one administration of FITC, whereas injection of *Nr4a3^+/+^* BMDCs without FITC treatment did not cause such swelling. We next s.c. injected FITC-treated *Nr4a3^+/+^* or *Nr4a3^-/-^* BMDCs into *Nr4a3^+/+^* or *Nr4a3^-/-^* mice and found that the transfer of FITC-treated *Nr4a3^+/+^* BMDCs caused ear swelling in both *Nr4a3^+/+^* and *Nr4a3^-/-^* mice, whereas the degree of ear swelling in *Nr4a3^+/+^* and *Nr4a3^-/-^* mice injected with FITC-treated *Nr4a3^-/-^* BMDCs was low (**Fig. 2c**). In addition, when CD11c^+^ cells that were isolated from the cLN of FITC-treated *Nr4a3^+/+^* mice were s.c. preinjected, the *Nr4a3^-/-^* and *Nr4a3^+/+^* mice exhibited similar levels of ear swelling (**Fig. 2d**).

These results suggest that NR4A3 deficiency diminishes DC migratory function, resulting in the suppression of CHS pathology accompanied by a decrease in T cells in the cLN.

### Localization of LCs and DCs in steady-state *Nr4a3*^-/-^ mice and expression of CCR7 in *Nr4a3*^-/-^ DCs

To evaluate the effects of NR4A3 deficiency on DCs, we investigated the localization of LCs and DCs in the skin of *Nr4a3^-/-^*mice in the steady state. Immunohistochemistry of the epidermal sheet (**Fig. 3a**) and flow cytometry of skin leukocytes (**Fig. 3b**) demonstrated that there were no significant differences between *Nr4a3^+/+^* and *Nr4a3^-/-^* in numbers of LCs stained with anti-MHCII antibody in the epidermal sheet (**Fig. 3a**), the percentages of MHCII^+^/CD11c^+^/Langerin^+^/EpCAM^+^ LCs in the epidermis or the percentages of MHCII^+^/CD11c^+^ dermal DCs in the dermis (**Fig. 3b**). The localization of DC subsets in the cLN in the steady state was analyzed by flow cytometry. As shown in **Fig. 3c**, the percentage (top) and the absolute number (bottom) of migDCs but not resDCs in the cLN was significantly reduced in the *Nr4a3^-/-^* mice. The percentage of MHCII^+^/CD11c^+^/Langerin^high^ LCs in the cLN was also markedly reduced in steady-state *Nr4a3^-/-^* mice (**Supplementary Fig. S2a**).

**Figure 3.**
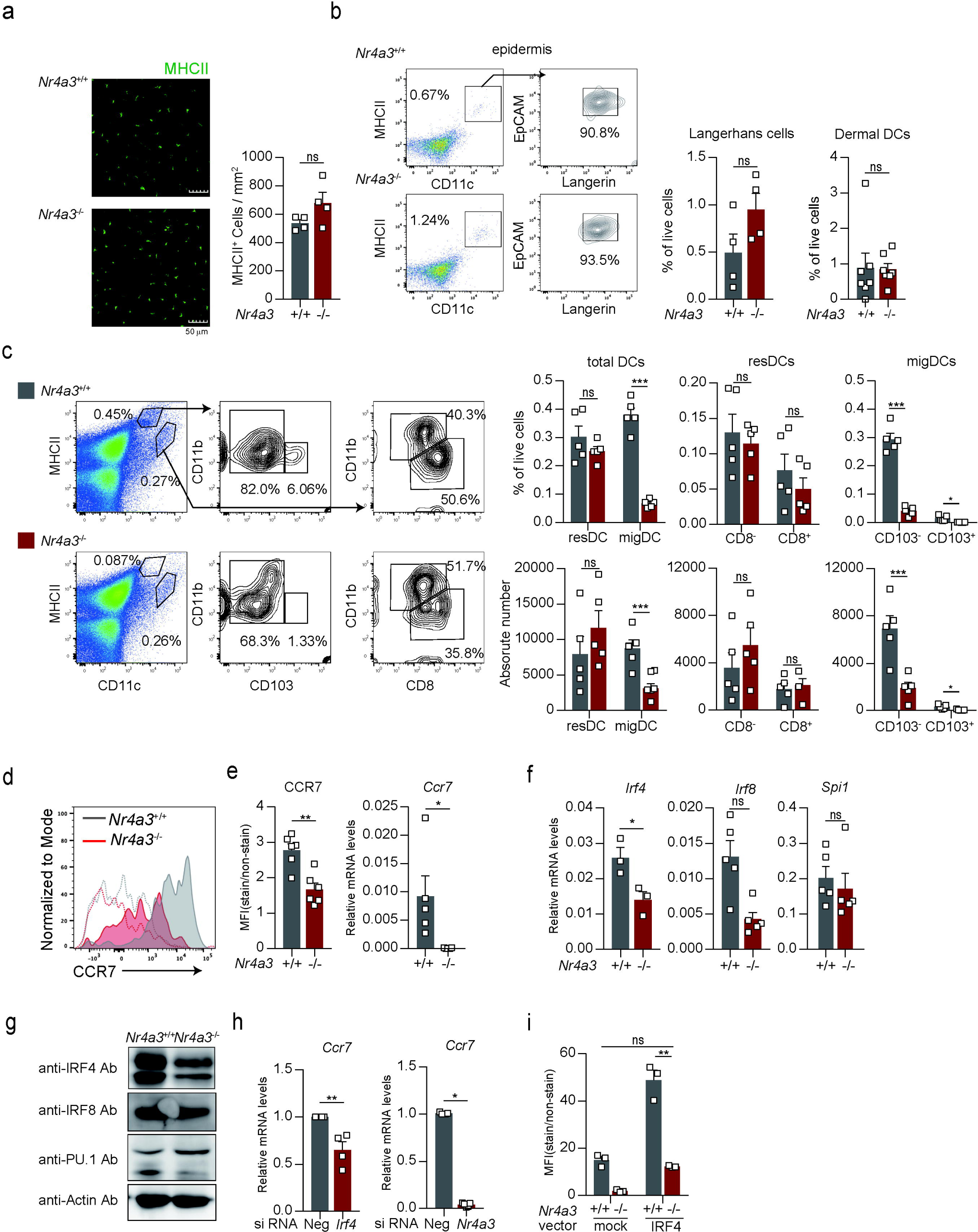
Effects of NR4A3 deficiency on the localization of DCs in disease-free mice and on the expression of CCR7 and related transcription factors in DCs. **a.** LCs in epidermal sheets. Left: FITC-labeled Ab staining for MHC class II on epidermal sheets from *Nr4a3^+/+^* mice (top) and *Nr4a3^-/-^* mice (bottom). Right: Absolute numbers of MHC class II-positive cells per unit area on epidermal sheets. **b.** Identification of epidermal LCs and dermal DCs by flow cytometry. Left: Gating strategy for epidermal LCs. Right: Proportions of epidermal LCs (MHCII^+^/CD11c^+^/EpCAM^+^/CD207^+^) and dermal DCs (MHCII^+^/CD11c^+^) among total live cells isolated from the epidermis and dermis, respectively. **c.** Identification of migDCs and resDCs in the cLNs. Left: Gating strategy for DCs in the cLNs. MHCII^high^/CD11c^int^ cells and MHCII^int^/CD11c^high^ cells were considered to be migDCs and resDCs, respectively. Right: Proportions (top) and absolute numbers (bottom) of each subtype of DCs in the cLNs. **d.** CCR7 expression levels on migDCs in the cLNs of *Nr4a3^+/+^* and *Nr4a3^-/-^* mice. MigDCs were identified by the same gating strategy, as shown in Fig. 3c. **e.** Expression levels of CCR7 in *Nr4a3^+/+^* and *Nr4a3^-/-^* BMDCs. Left: Cell surface protein levels of CCR7. Right: mRNA levels of *Ccr7*. **f.** Messenger RNA levels of *Irf4*, *Irf8* and *Spi1* in *Nr4a3^+/+^* and *Nr4a3^-/-^* BMDCs. **g.** Western blot profiles showing the protein expression levels of IRF4, IRF8, and PU.1 in whole-cell lysates of *Nr4a3^+/+^* and *Nr4a3^-/-^* BMDCs. Lysate aliquots containing 15 μg of protein were loaded into each lane. **h.** *Ccr7* mRNA levels in siRNA-introduced BMDCs. BMDCs generated from control mice were transfected with *Irf4* siRNA (left), *Nr4a3* siRNA (right), or negative controls. BMDCs were collected to measure mRNA levels at 24 h and 48 h after transfection with *Irf4* siRNA and *Nr4a3* siRNA, respectively. The knockdown degrees of the *Irf4* and *Nr4a3* mRNAs are shown in **Supplementary Fig. S2b**. **i.** Cell surface expression of CCR7 on IRF4-overexpressing DCs. *Nr4a3^+/+^*and *Nr4a3^-/-^* BMDCs were transfected with pIRES2-AcGFP1-Myc-IRF4 (IRF4) or its control pIRES2-AcGFP1 (mock). BMDCs were stimulated with 100 ng/mL of LPS at 24 h after plasmid introduction. After an additional 24 h of incubation, the BMDCs were harvested and analyzed via flow cytometry. The MFIs of CCR7 in the transfectants (GFP-positive cells) were measured. All of the results in Fig. 3 are shown as the mean ± SE. Significance was determined by two-tailed unpaired Student’s *t*-test (**a**, **b**, **c**, **e, f, h**), or one-way ANOVA with the Tukey-Kramer multiple comparison test (**g**). \**p*<0.05; \*\**p*<0.01; \*\*\**p*<0.001; ns, not significant.

A previous valuable study of *Nr4a3*-deficient mice demonstrated a reduced number of CD103^+^ migDCs in the MLN due to a defect in CCR7 expression on *Nr4a3*-deficient DCs ^11^. Under our experimental conditions, we also found that the expression levels of CCR7 on migDCs in the cLNs of steady-state *Nr4a3*^-/-^ mice (**Fig. 3d**) and on BMDCs generated from *Nr4a3*^-/-^ mice (**Fig. 3e, left**) were lower than those in *Nr4a3*^+/+^ mice, which was consistent with the reduced expression of *Ccr7* mRNA in *Nr4a3^-/-^* DCs (**Fig. 3e, right**).

The transcription factor FOXO1, which was decreased in *Nr4a3*^-/-^ DCs, was previously identified to cause the reduced expression of CCR7 on CD103^+^ DCs ^11^. In the present study, we focused on the hematopoietic cell-specific transcription factor IRF4 based on previous observations showing that the expression level of IRF4 in BMDCs was decreased by transfection with *Nr4a3* siRNA ^10^, and that cutaneous DCs in *Irf4*^-/-^ mice exhibit impaired migration to lymph nodes due to a lack of CCR7 expression ^13^. Quantitative PCR and western blotting analyses revealed that the mRNA level of *Irf4* in BMDCs was significantly reduced in *Nr4a3^-/-^* DCs (**Fig. 3f**), which was reflected in the protein level of IRF4 (**Fig. 3g**), whereas the protein level of PU.1 was increased by NR4A3 deficiency. We also revealed that knockdown (KD) of IRF4 and NR4A3 via siRNA transfection significantly decreased *Ccr7* mRNA levels in BMDCs (**Fig. 3h**), and exogenous overexpression of IRF4 restored the cell surface expression of CCR7 in *Nr4a3*^-/-^ BMDCs and enhanced CCR7 expression in *Nr4a3*^+/+^ BMDCs (**Fig. 3i**).

These results indicate that reduced expression of IRF4 is a cause of low expression of CCR7 in *Nr4a3*^-/-^ DCs, resulting in a small number of migDCs in the cLNs of *Nr4a3*^-/-^ mice.

### Effects of NR4A3 deficiency on psoriasis

We next utilized an imiquimod-induced psoriasis model and found that the degree of swelling in the imiquimod-treated ear of *Nr4a3*^-/-^ mice was significantly greater than that in the treated ear of *Nr4a3*^+/+^ mice, particularly during the later period (Days 4-7) (**Fig. 4a**). When mRNA levels of cytokines in the ear skin were measured on Day 7, significantly increased expression of inflammatory cytokines, including *Il6*, *Il12b*, *Il17a*, *Il17f*, and *Il23a*, was detected in the ears of imiquimod-treated *Nr4a3*^-/-^ mice (**Fig. 4b**). We also found that the number of γδT cells in the lesioned skin was significantly increased in the *Nr4a3*^-/-^ mice (**Fig. 4c**) and that the localization of Th1 and Th17 cells in the draining cLNs of the imiquimod-treated ear was reduced in *Nr4a3*^-/-^ mice (**Fig. 4d**).

**Figure 4.**
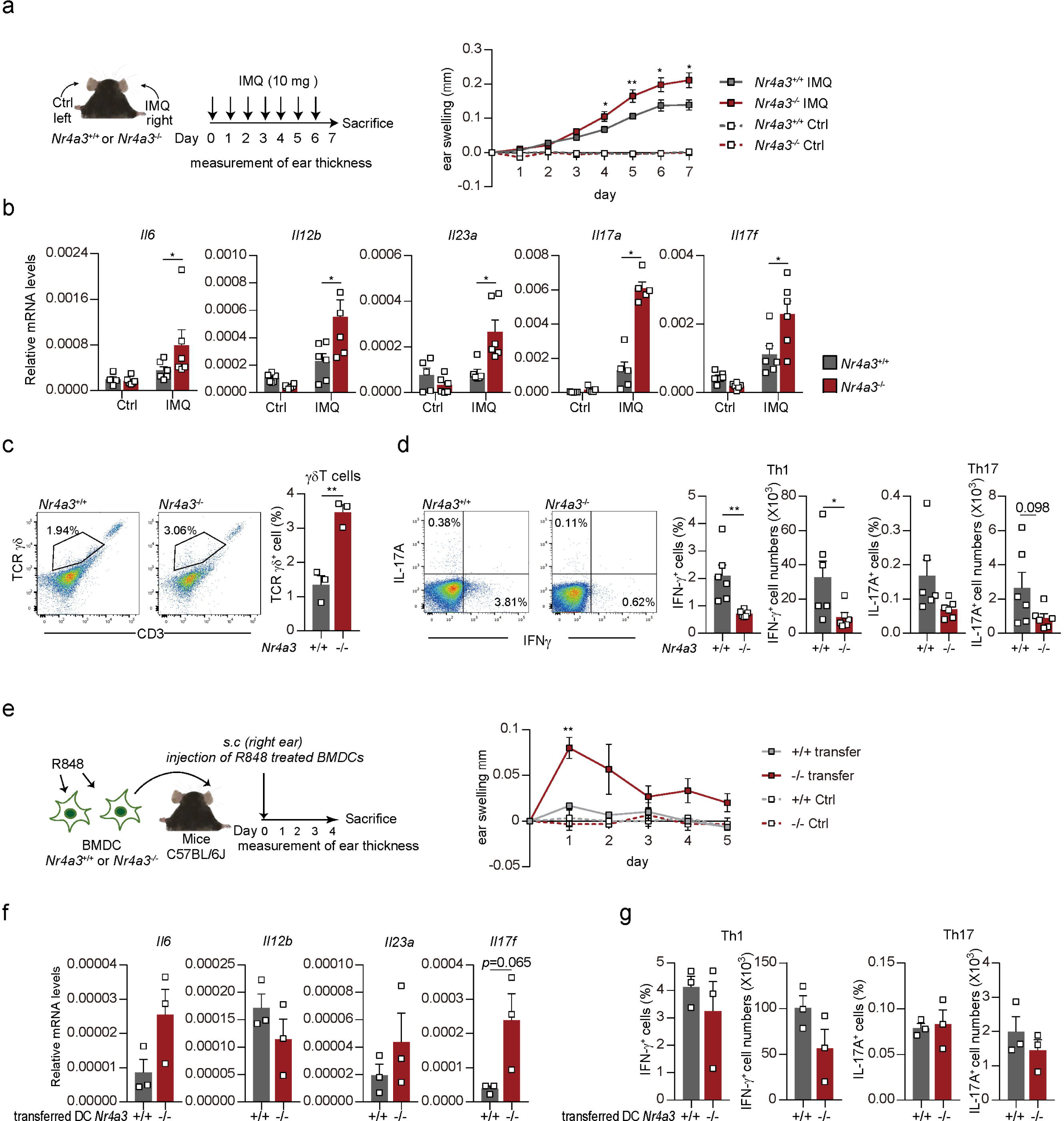
Pathology of psoriasis in NR4A3-deficient mice and effects of NR4A3 deficiency on the function of TLR7 ligand-stimulated DCs in vivo. **a.** Schematic of imiquimod (IMQ)-induced psoriasis model (left). The right ears of ten-week-old male *Nr4a3^+/+-^* and *Nr4a3^-/-^*mice were topically administered 10 mg of BESELNA cream. From Day 0 to Day 7, the ear thickness of the psoriasis model mice was assessed daily. Ear swelling = (ear thickness on each day) – (ear thickness on Day 0). **b.** Messenger RNA levels of cytokines in the ear skin of psoriasis-induced mice on Day 7. Ctrl, the left ear without IMQ; IMQ, the right ear with IMQ-treatment. **c.** Number of γδT cells in psoriasis skin lesions. **d.** IFN-γ-positive or IL-17A-positive CD4^+^ T cells in the cLNs of psoriasis mice. **e.** R848-stimulated *Nr4a3^+/+^* or *Nr4a3^-/-^* BMDCs were transferred into control C57BL/6 mice. Left: Schematic of the experimental procedure. *Nr4a3^+/+^* or *Nr4a3^-/-^* BMDCs were incubated in the presence of 1 μg/mL R848 in culture media for 1 h. After washing with PBS two times, 6.25 x 10^5^ cells of *Nr4a3^+/+^* or *Nr4a3^-/-^* BMDCs treated with R848 were s.c. injected into the right ear of C57BL/6 mice on Day 0. Right: Ear swelling determined in the same way as in Fig. 4a. **f.** Messenger RNA levels of cytokines in the ear skin of BMDC-transferred mice on Day 5. **g.** IFN-γ-positive or IL-17A-positive CD4^+^ T cells in the cLNs of BMDC-transferred mice. All of the results in Fig. 4 are shown as the mean ± SE. Significance was determined by two-tailed unpaired Student’s *t* test (**c**, **d, f**, **g**), or one-way ANOVA with the Tukey-Kramer multiple comparison test (**a**, **b, e**). \**p*<0.05; \*\**p*<0.01; \*\*\**p*<0.001; \*\*\*\**p*<0.0001; ns, not significant.

### R848-treated *Nr4a3*^-/-^ DCs induced inflammation in the skin

To reveal the effects of NR4A3 deficiency in DCs on psoriasis, we injected R848-primed BMDCs generated from *Nr4a3*^+/+^ or *Nr4a3*^-/-^ mice into control C57BL/6 mice. As shown in **Fig. 4e**, the ear thickness of mice s.c. injected *Nr4a3*^-/-^ BMDCs was significantly increased compared with that of mice injected with *Nr4a3*^+/+^ BMDCs. Under these experimental conditions, increased expression levels of *Il6*, *Il17f*, and *Il23a* mRNAs in the lesioned skin (**Fig. 4f**) and slightly reduced numbers of Th1 and Th17 cells in the cLNs (**Fig. 4g**) were observed in *Nr4a3*^-/-^ BMDC-injected mice.

These results suggest that NR4A3-deficient DCs amplify TLR7-dependent inflammation in the skin.

### Roles of NR4A3 in the expression of TLR7 in DCs

We investigated the TLR-mediated activation of *Nr4a3*^-/-^ DCs to clarify the effects of NR4A3 knockout (KO) on the gene expression and function of DCs. When CD11c^+^ DCs isolated from the spleen were stimulated with LPS, no apparent differences in the amounts of IL-6, IL-12p40, or TNF-α released from the stimulated DCs were observed between the *Nr4a3*^+/+^ and *Nr4a3*^-/-^ (**Fig. 5a**). In contrast, the release of cytokines upon R848 stimulation was significantly increased in *Nr4a3*^-/-^ DCs (**Fig. 5a**). To reveal the mechanism by which TLR7-dependent stimulation of DCs was amplified by NR4A3 deficiency, we compared *Tlr7* mRNA levels between *Nr4a3*^+/+^ and *Nr4a3*^-/-^ DCs. As shown in **Fig. 5b**, R848 treatment upregulated *Tlr7* mRNA expression in DCs, which was significantly increased in NR4A3-deficient DCs, whereas *Tlr4* mRNA expression was not affected by NR4A3 deficiency (**Fig. 5b**). Furthermore, the *Tlr7* mRNA level in psoriatic skin lesions was significantly higher in *Nr4a3*^-/-^ mice than in *Nr4a3*^+/+^ mice (**Fig. 5c**).

**Figure 5.**
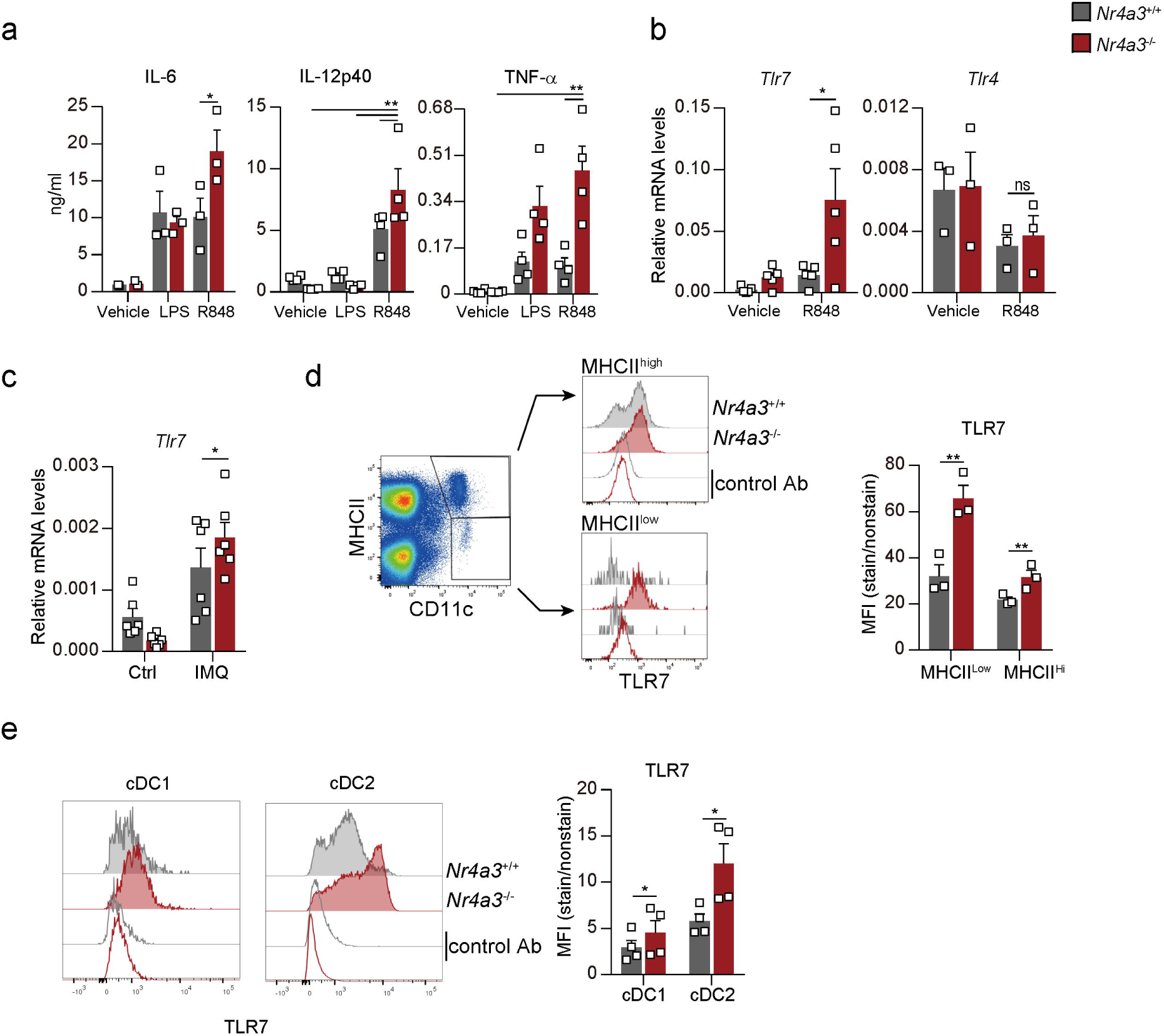
Enhancement of TLR7-mediated cytokine production and TLR7 expression in NR4A3 knockout DCs. **a.** Cytokine production by splenic DCs. CD11c^+^ cells (2 × 10^5^) isolated from the spleens of *Nr4a3^+/+^* and *Nr4a3^-/-^* mice were incubated in the presence of 100 ng/mL LPS, 1 μg/mL R848, or vehicle (finally 0.1% H_2_O: DMSO = 1:1) for 24 h. Concentrations of IL-6, IL-12p40, and TNF-α in culture supernatants were determined via ELISA. **b.** Messenger RNA levels of TLRs in BMDCs. *Nr4a3^+/+^* and *Nr4a3^-/-^* BMDCs were cultured in the presence of 1 μg/mL R848, or vehicle (0.1% DMSO) for 4 h, and were subsequently collected to determine the mRNA levels of *Tlr4* and *Tlr7*. **c.** *Tlr7* mRNA levels in the skin. On Day 7 of the IMQ-induced psoriasis model, the ears of the *Nr4a3^+/+^* and *Nr4a3^-/-^* mice were collected just after scarifice. Ctrl, left ear without any applications; IMQ, right ear applied BESELNA cream. **d.** Protein expression of TLR7 in splenic DCs. Whole-spleen cells from *Nr4a3^+/+^* and *Nr4a3^-/-^* mice were stained with anti-I-A/I-E, anti-CD11c, and anti-TLR7 Abs. **e.** Protein expression of TLR7 in cDC1s and cDC2s. Conventional DC1s and cDC2s in the spleen were characterized with the gating strategies shown in **Supplementary Fig. S3**. All of the results in Fig. 5 are shown as the mean ± SE. Significance was determined by two-tailed unpaired Student’s *t* test (**a**, **d**, **e**), or one-way ANOVA with the Tukey-Kramer multiple comparison test (**b**, **c**). \**p*<0.05; \*\**p*<0.01; ns, not significant.

We then examined the protein expression levels of TLR7 in splenic CD11c^+^ DCs by flow cytometry and found that *Nr4a3*^-/-^ DCs expressed significantly higher levels of TLR7 than *Nr4a3*^+/+^ DCs among both MHC II^high^ DCs and MHC II^low^ DCs (**Fig. 5d**). When splenic cDCs, distinguished as cDC1s and cDC2s, were analyzed, increased expression of TLR7 was observed in both *Nr4a3*^-/-^ cDC2s and cDC1s, and this change was more striking in cDC2s (**Fig. 5e**).

These results indicate that *Nr4a3*^-/-^ DCs released higher amounts of inflammatory cytokines upon TLR7-mediated stimulation due to the elevated mRNA and protein levels of TLR7 induced by NR4A3 deficiency.

### Effects of NR4A3 deficiency on the development and gene expression of DCs

The abovementioned results in the present study revealed that NR4A3 deficiency decreased the migratory activity and increased the TLR7 signaling response of DCs, which are involved in relieving the pathology of CHS and exacerbating psoriasis, respectively. To further clarify the effects of NR4A3 deficiency on the development and gene expression of DCs, we compared *Nr4a3*^+/+^ BMDCs and *Nr4a3*^-/-^ BMDCs.

Flow cytometry analysis revealed that the proportion of CD86^high^/MHCII^high^ cells was reduced in *Nr4a3*^-/-^ BMDCs (**Fig. 6a**), suggesting that there was a delay or insufficiency in the GM-CSF-induced development of DCs. In addition, the mRNA levels of *Irf4* and *Irf8* but not *Spi1* were significantly decreased in *Nr4a3*^-/-^ BMDCs even under stimulated conditions (**Fig. 6b**). When NR4A3 was knocked down in mature BMDCs, a decrease in the IRF4 protein level was still observed under stimulated conditions (**Fig. 6c**). Taken together with these results and those shown in **Fig. 3f**, and **3g**, we conclude that NR4A3 is required for the expression of IRF4 in nonstimulated and stimulated DCs. Among the heterozygous populations of GM-CSF-induced BMDCs, the MHCII^high^ population is characterized by DC-like cells, and the MHCII^int^ population comprises macrophage-like cells^14^. In our previous study, knocking down IRF4 or PU.1 via siRNA transfection suppressed the development of DC-like cells and the DC-like cell-specific expression of PD-L2 in BMDCs^15^. To confirm the involvement of reduced expression of IRF4 in delaying or suppressing the development of the MHCII^high^ population in *Nr4a3*^-/-^ BMDCs, we overexpressed IRF4 in *Nr4a3*^-/-^ BMDCs by using a retroviral vector. As shown in **Fig. 6d**, exogenous expression of IRF4 abrogated the decrease in the proportion of the MHCII^high^ population.

**Figure 6.**
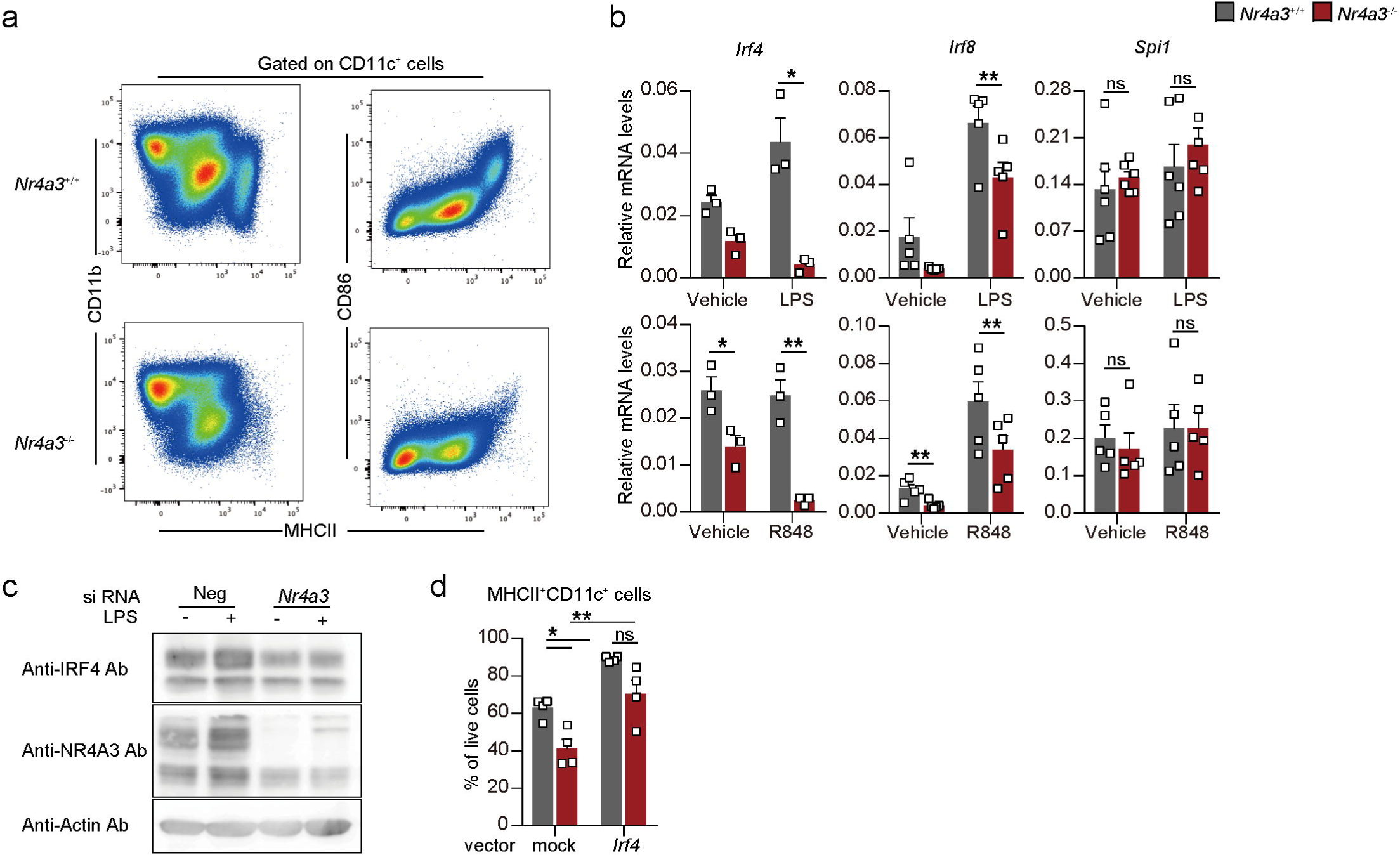
Effects of NR4A3 deficiency on the development and transcriptional regulation of DCs. **a.** Flow cytometric analysis of BMDCs. Cell surface expression of MHCII, CD86, and CD11b on *Nr4a3^+/+^* and *Nr4a3^-/-^* BMDCs, which were pregated as CD11c^+^ cells. **b.** Messenger RNA levels of transcription factors in BMDCs with or without stimulation. *Nr4a3^+/+^* and *Nr4a3^-/-^* BMDCs were incubated in the presence of 100 ng/mL LPS, 1 μg/mL R848, or vehicle (H_2_O for LPS, DMSO for R848) for 4 h. **c.** Protein expression levels of transcription factors in siRNA-introduced BMDCs. BMDCs generated from control BALB/c mice were transfected with *Nr4a3* siRNA or the corresponding control. After 24 h of electroporation, the BMDCs were stimulated with 100 ng/mL LPS and cultivated for an additional 24 h. **d.** Effect of IRF4 overexpression on the proportion of the MHCII^+^/CD11c^+^ population in BMDCs. *Nr4a3^+/+^* and *Nr4a3^-/-^*BMDCs were transfected with retrovirus carrying IRF4 cDNA and dsRed cDNA linked to the with IRES sequence (*Irf4*) or with a control vector containing dsRed cDNA (mock). The frequency of MHCII^+^/CD11c^+^ cells among the dsRed-positive cells is shown. All of the results in Fig. 6b are shown as the mean ± SE. Significance was determined by two-tailed unpaired Student’s *t*-test. \**p*<0.05; \*\**p*<0.01; ns, not significant.

### PU.1 is involved in the enhanced expression of IRF7 in *Nr4a3*^-/-^ DCs

We further investigated the expression levels of IRF2, 3, 5, and 7, which are transcription factors involved in TLR7-mediated gene expression in DCs^16^. Quantitative PCR revealed that stimulation by LPS or R848 upregulated *Irf7* mRNA levels in BMDCs and that the R848-induced increase in *Irf7* mRNA levels was significantly greater in *Nr4a3*^-/-^ BMDCs than in control BMDCs, whereas the mRNA levels of *Irf2*, *Irf3*, and *Irf5* were not affected by stimulation or NR4A3 deficiency (**Fig. 7a**). Considering the important role of IRF7 in the expression of type I IFNs in DCs^17^, we examined the mRNA levels of *Ifna4* and *Ifnb1* to confirm whether the expression of type I IFNs was increased in *Nr4a3*^-/-^ BMDCs. As shown in **Fig. 7b (left)**, the expression of *Ifna4* mRNA was induced by LPS stimulation and was significantly increased in *Nr4a3*^-/-^ BMDCs. Furthermore, transcription of the *Ifnb1* gene was induced by not only LPS but also R848, and stimulation-induced increases in *Ifnb1* mRNA levels were markedly enhanced by the NR4A3 deficiency (**Fig. 7b right**). We also revealed that upregulation of IRF7 and type I IFNs by NR4A3 KO, which was caused in GM-CSF-induced BMDCs (**Fig. 7a** and **b**), was not observed in Flt3L-induced BMDCs (**Supplementary Fig. S4**). When we analyzed the binding profiles of transcription factors to the *Irf7* gene using the ChIP-Atlas database, we found specific binding of PU.1 to the intron of the *Irf7* gene in both cDC and pDC, whereas apparent binding of PU.1 was not observed for the *Irf3* gene (**Fig. 7c**). Considering that KO of NR4A3 tended to increase PU.1 protein levels in DCs (**Fig. 3g**), we hypothesized that *Irf7* upregulation and subsequently upregulation of *Ifnb1* occurred through the enhanced activity of PU.1. To reveal the involvement of PU.1 in the mRNA expression of *Irf7* and *Ifnb1*, we examined the effect of PU.1 KD on the expression of *Irf7* and *Ifnb1*. As expected, the mRNA levels of *Irf7* and *Ifnb1* were significantly decreased in *Spi1* siRNA-introduced DCs, whereas those of *Irf3* and *Ifna4* were unaffected by PU.1 KD (**Fig. 7d**). The transcript level of *Ccr7* was also markedly decreased in PU.1 KD DCs, which was likely a result of the decreased expression of the *Irf4*, which is under the control of PU.1 ^18^, and to the KD of PU.1 itself, which transactivates the *Ccr7* gene in DCs ^19^.

**Figure 7.**
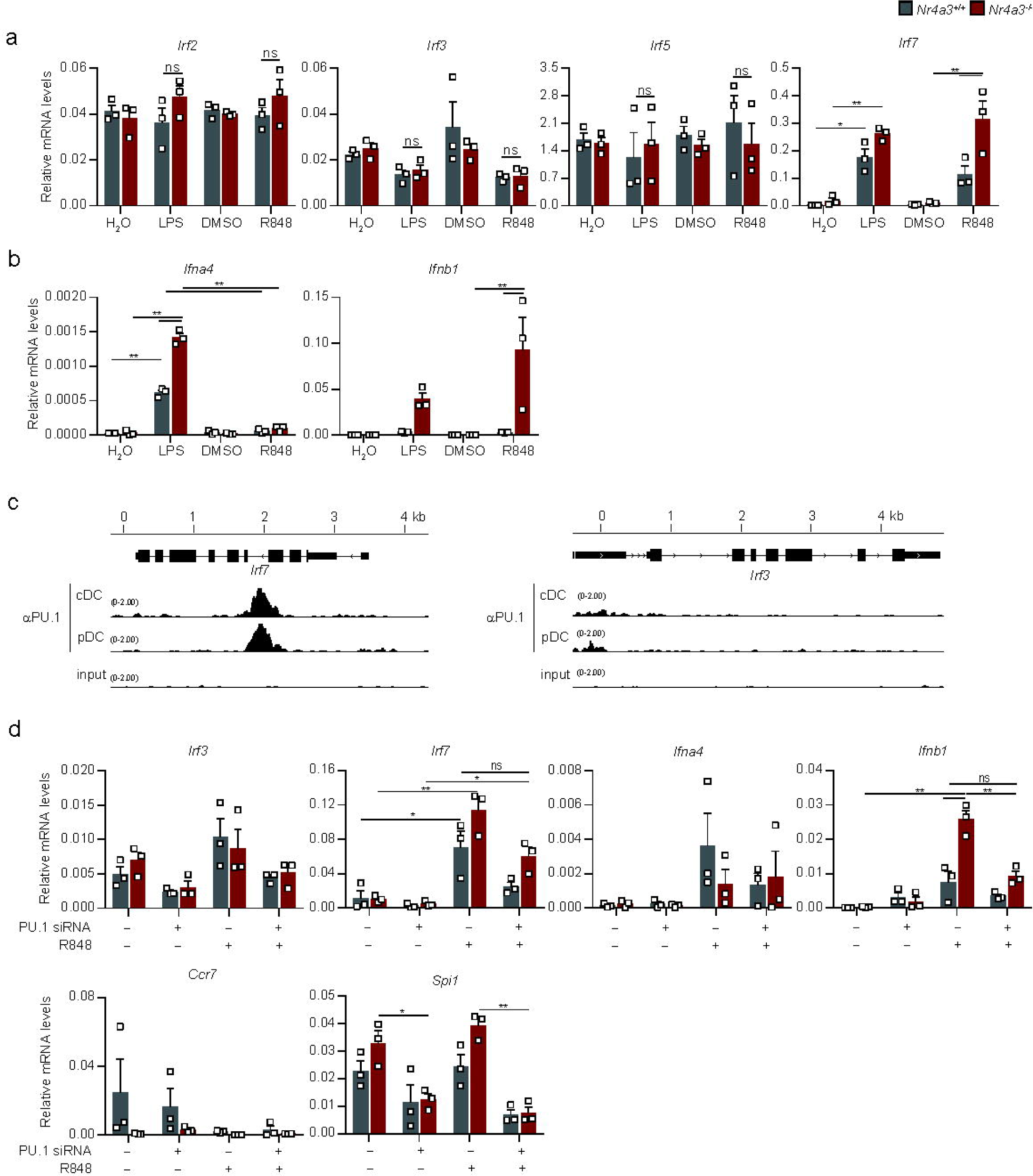
Involvement of NR4A3 in the expression of IRF family molecules. **a.** Messenger RNA levels of *Irf2*, *Irf3*, *Irf5*, and *Irf7* in BMDCs. **b.** Messenger RNA levels of *Ifna4* and *Ifnb1* in BMDCs. *Nr4a3^+/+^* and *Nr4a3^-/-^* BMDCs were incubated in the presence of 100 ng/mL LPS, 1 μg/mL R848, or vehicle (0.1% H_2_O for LPS, and DMSO for R848) for 24 h (**a** and **b**). **c.** PU.1 binding profiles of the *Irf7* and *Irf3* genes in DCs. ChIP-seq tracks for PU.1 binding within the *Irf7* (left) and *Irf3* (right) loci in cDCs (GM-CSF-induced BMDCs) and pDCs (Flt3L-induced BMDCs) obtained by recalculating ChIP-Atlas data (cDCs, https://chip-atlas.org/view?id=SRX4909225; pDCs, https://chip-atlas.org/view?id=SRX4909226; input, https://chip-atlas.org/view?id=SRX4909224). **d.** Messenger RNA levels in PU.1 KD DCs. BMDCs generated from control C57BL/6 mice were transfected with *Spi1* siRNA or the corresponding control. At 24 h after siRNA transfection, 100 ng/mL LPS or 1 μg/mL R848 was added to the culture media. After an additional 24 h of incubation, the BMDCs were collected to measure the mRNA levels. All of the results in Fig. 7 are shown as the mean ± SE. Significance was determined by ANOVA with the Tukey-Kramer multiple comparison test. \**p*<0.05; \*\**p*<0.01; ns, not significant.

Taken together, these results indicate that NR4A3 is involved in regulating the expression of IRF7 by controlling the expression and/or function of PU.1 in DCs, which affects TLR-mediated cytokine gene expression.

## Discussion

NR4A3 is a ligand-independent transcription factor that belongs to the nuclear receptor superfamily. The involvement of NR4A3 in several biological processes, including the development of the central nervous system^2,3^, pathogenesis of atherosclerosis^5,6^, and defense against bacterial infection^11^ has been demonstrated by studies using *Nr4a3*^-/-^ mice. In the present study, we revealed the opposite effects of NR4A3 deficiency on CHS and psoriasis in mice. Specifically, *Nr4a3*^-/-^ mice were resistant to CHS and sensitive to psoriasis. Dysfunction in DCs (i.e., decreased expression of CCR7, enhanced expression and function of TLR7, and dysregulation of the transcription factors IRF4, IRF7, and PU.1 in DCs) were identified as factors, which were associated with the amelioration of CHS and exacerbation of psoriasis. Although NR4A3 is a ubiquitous molecule, we have shown that DC dysfunction in *Nr4a3*^-/-^ mice is an important factor modulating the pathogenesis of these skin diseases (**Fig. 8**).

**Figure 8.**
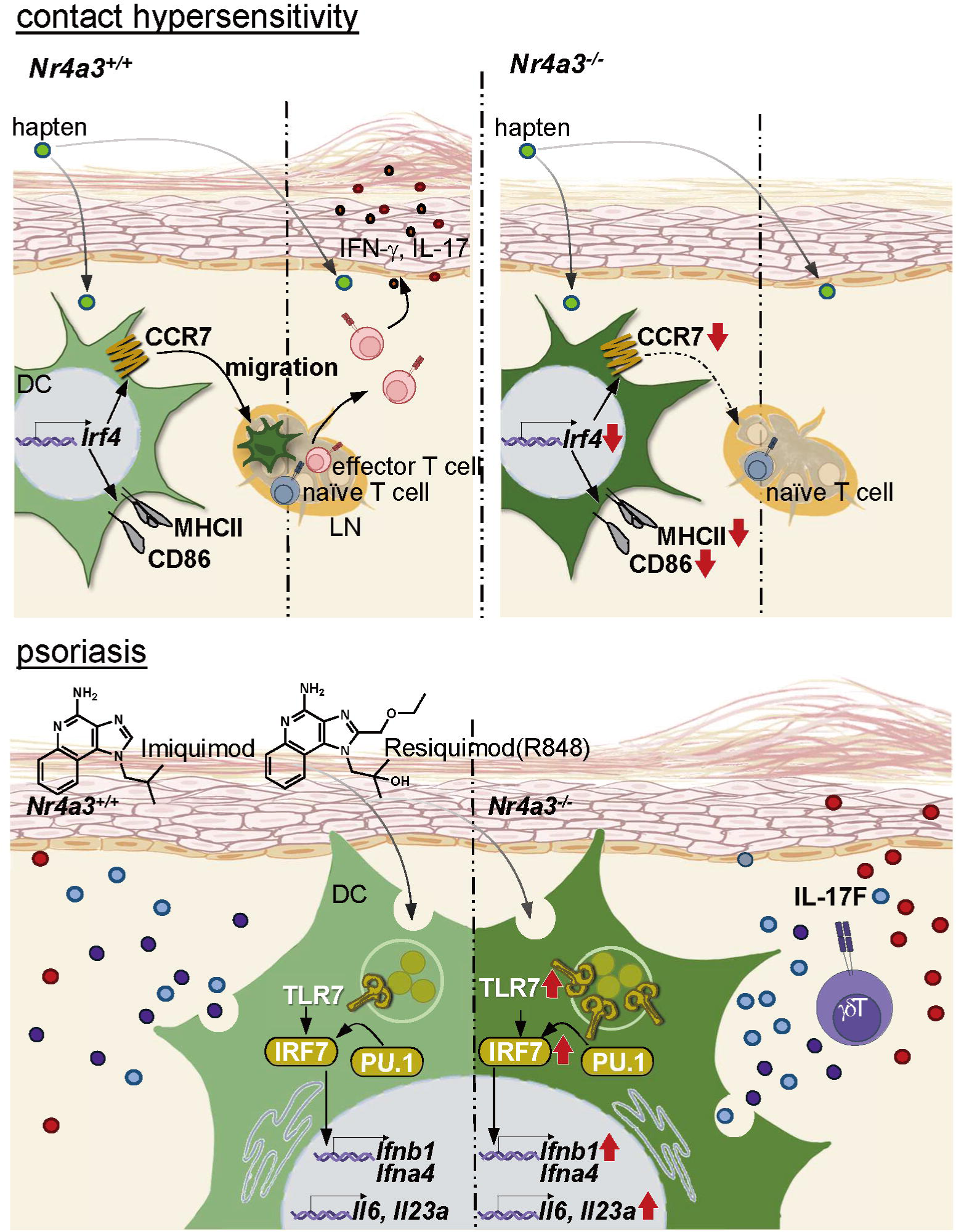
Schematic drawing showing that *Nr4a3*^-/-^ DC dysfunction ameliorates CHS and exacerbates psoriasis in mice.

We demonstrated that the surface expression of CCR7 on DCs was markedly reduced by NR4A3 deficiency, which decreased the migratory activity of DCs, resulting in the alleviation of CHS. A reduction in CCR7 levels on CD103^+^ DCs in *Nr4a3*^-/-^ mice was reported in a previous study^11^, in which FOXO1 was identified as a transcription factor regulating the expression of the *Ccr7* gene under the control of NR4A3. In contrast, the present study suggested that reduced levels of IRF4 are involved in the downregulation of CCR7 expression in *Nr4a3*^-/-^ DCs. A previous finding showed that the failure of dermal DCs to migrate to the LN and a reduction of surface CCR7 in *Irf4*^-/-^ mice induced CHS^13^, which is consistent with our hypothesis that the NR4A3-IRF4-CCR7 axis regulates DC migration and the development of CHS. In a study showing that NR4A3 is required for surface CCR7 expression and cross-presentation in monocyte-derived DCs (MoDCs), IRF4 expression in *Nr4a3*^+/+^ and *Nr4a3*^-/-^ MoDCs was equivalent ^12^. However, the reduced expression of CCR7 on *Irf4*^-/-^ MoDCs was restored by exogenous overexpression of NR4A3^12^, indicating the existence of an IRF4-NR4A3-CCR7 axis. We cannot explain this discrepancy in the order of gene regulation; however, the regulatory relationship between IRF4 and NR4A3 may be due to the fact that NR4A3 and IRF4 transactivate the *Ccr7* gene while regulating the expression of each other. Alternatively, this difference may be due to the DC subset, since IRF4 is crucial for the development and gene expression of cDC2s^20^. That is, IRF4 is required for *Ccr7* expression in cDC2s, whereas NR4A3 is essential for *Ccr7* expression in MoDCs.

Increased expression of the TLR7 protein and *Tlr7* mRNA induced by NR4A3 deficiency was observed in cDCs, particularly in cDC2s. Although human pDCs, which produce IFN-α and play an important role in psoriasis ^21^, are known to constitutively express TLR7 ^22^, TLR7 expression on pDCs and *Ifna4* mRNA levels in Flt3L-induced BMDCs were not affected by a lack of NR4A3 (**Supplementary Fig. S4**). The marked increase in *Ifnb1* mRNA levels in GM-CSF-induced *Nr4a3*^-/-^ BMDCs, which can be produced by cDCs, compared to the lower magnitude of increase in *Ifna4* mRNA, which can be produced by pDCs but not by cDCs ^23,24^, also suggested that the effects of NR4A3 KO are limited to cDCs (especially cDC2s) among DC subsets. A previous observation that selective TLR7-mediated activation of Langerin-negative cDCs is sufficient to induce psoriasis plaque formation through the production of IL-23, independent of pDCs or type I IFNs ^25^, might support our hypothesis that the hyperresponse of *Nr4a3*^-/-^ cDCs to TLR7 ligands is critical for exacerbating psoriasis in *Nr4a3*^-/-^ mice. However, whether NR4A3 deficiency affects the production levels of type I IFNs in steady-state and psoriasis-induced mice has not yet been investigated. Further detailed analysis is still needed to clarify the roles of NR4A3 in TLR7-mediated signaling and the production of type I IFNs. PU.1 is a hematopoietic cell-specific transcription factor that belongs to the Ets-family. The essential roles of PU.1 in the development and gene expression of DCs are well known ^26–28^. PU.1 transactivates target genes by binding to the Ets-motif alone or to the EICE-motif with forming a heterodimeric complex with IRF4 or IRF8. Although the binding of PU.1 to the *Irf7* gene in Flt3L cultured pDCs has been previously shown ^29^, the role of PU.1 in the transcription of the *Irf7* gene has not been determined. We found that PU.1 binds to the *Irf7* gene not only in pDCs but also in cDCs via in silico analysis and that PU.1 KD significantly decreased *Irf7* mRNA levels in GM-CSF-induced BMDCs. *Irf7* mRNA levels were increased in *Nr4a3*^-/-^ BMDCs, in which the protein levels of IRF4 and IRF8 were decreased markedly and moderately, respectively, indicating that PU.1 positively regulates the expression of the *Irf7* gene without forming a heterodimer with IRF4 or IRF8. The *Ifnb1* mRNA level was also decreased by PU.1 KD. IRF7 and IFN-β affect each other’s the gene expression. That is, IRF7 is crucial for the transcription of the *Ifnb1* gene, and IFN-β signaling promotes the transcription of the *Irf7* gene ^30^. Although it has not been determined whether the KD of PU.1 directly suppressed *Ifnb1* transcription or indirectly through the downregulation of *Irf7* transcription, our KD experiments suggest that PU.1 is required for the expression of both *Irf7* and *Ifnb1*.

Here, we identified IRF4 and PU.1 as transcription factors associated with the dysregulation of the *Ccr7* and *Irf7*/*Ifnb* genes in *Nr4a3*^-/-^ DCs. However, the mechanism underlying the enhanced expression and function of TLR7, which is involved in increased inflammation in psoriasis, is unclear at present. Although we cannot exclude the possibility that NR4A3 directly regulates the *Tlr7* gene, the suppressive effect of NR4A3 on the expression of TLR7 may reflect a negative feedback function of NR4A3 that is rapidly activated upon various stimuli.

DCs control innate and adaptive immune responses through various processes, including the release of inflammatory cytokines, and the activation of naïve T cells via its antigen presenting activity. Obviously, it is important and interesting to examine the pathogenesis of various immune-related diseases in *Nr4a3*^-/-^ mice that exhibit DC dysfunctions.

## Methods

All mouse experiments were performed following the guidelines of the Institutional Review Board of Tokyo University of Science, and the present study was approved by the Animal Care and Use Committees of Tokyo University of Science: K22005, K21004, K20005, K19006, K18006, K17009, K17012, K16007, and K16010.

### Mice and cells

C57BL/6J mice were purchased from Japan SLC (Hamamatsu, Japan). *Nr4a3^-/-^*(*Nr4a1*^fl^/*Nr4a2*^fl^/*Nr4a3*^-/-^) and *Nr4a3^+/+^*(*Nr4a1*^fl^/*Nr4a2*^fl^/*Nr4a3*^+/+^) mice were generated in a previous study by Sekiya et al.^7^. Mice were maintained under specific pathogen-free-conditions. BMDCs were generated by cultivating BM cells supplemented with recombinant mouse GM-CSF (#576308, BioLegend) or recombinant mouse Flt3L (550706, BioLegend) as previously described ^15,27,31^. The cLNs were incubated in 100 μL of RPMI-1640 containing 10 mM HEPES (pH 7.4), 50 μg/mL Liberase TL, and 200 μg/mL DNaseI for 20 min at 37 ℃ with shaking to obtain a single-cell suspension. CD11c MicroBeads UltraPure, mouse (#130-125-835; Myltenyi Biotec) and a MojoSort Mouse CD4 T cell Isolation Kit (#480033) were used to collect CD11c^+^ DCs and CD4^+^ T cells from the spleen and/or LNs. Naïve CD4^+^ T cells isolated from the mouse spleen using the MojoSort Mouse Naïve CD4 T Cell Isolation Kit (#480040, BioLegend) were stimulated in a plate coated with LEAF purified anti-mouse CD3ε antibody (clone 145-2111C, BioLegend) and anti-CD28 antibody (clone 37.51, BioLegend) under Th1-polarizing condition (10 ng/mL recombinant mouse IL-12 (p70) (#577004, BioLegend), and 10 μg/mL anti-mouse IL-4 antibody (clone 11B11, BioLegend)) or Th2-polarizing condition (10 ng/mL recombinant mouse IL-4 (#574304, BioLegend), and 10 μg/mL anti-mouse IL-12/23p40 antibody (clone C17.8, BioLegend)).

### Reagents

DMSO (07-4860-5, Sigma Aldrich), LPS (L3024, Wako), and R848 (AG-CR1-3582-M005, AdipoGen) were purchased from the indicated sources.

The following antibodies were used in flowcytometry, which were obtained from indicated sources: APC anti-mouse I-A/I-E (MHC II) (clone M5/114.15.2, BioLegend, 1:500), FITC anti-mouse I-A/I-E (MHC II) (clone M5/114.15.2, BioLegend, 1:500), PerCP anti-mouse I-A/I-E (MHC II) (clone M5/114.15.2, BioLegend, 1:250), PerCP anti-mouse CD3ε (clone 145-2C11, BioLegend, 1:100), FITC anti-mouse CD4 (clone GK1.5, BioLegend, 1:400), PECy7 anti-mouse CD4 (clone GK1.5, Tonbo Biosciences, 1:1000), VioGreen anti-mouse CD8a (clone 53-6-7, Myltenyi Biotec, 1:20), APCCy7 anti-human/mouse CD11b (clone M1/70, Tonbo Biosciences, 1:2000), PECy7 anti-mouse CD11c (clone N418, Tonbo Biosciences, 1:1000), PerPC/Cy5.5 anti-mouse CD24 (clone M1/69, BioLegend, 1:500), FITC anti-human/mouse CD44 (clone IM7, BioLegend, 1:100), FITC anti-mouse/human CD45R/B220 (clone RA3-6B2, BioLegend, 1:100), PE anti-mouse CD62L (clone MEL-14, Tonbo Biosciences, 1:100), FITC anti-mouse CD86 (clone GL-1, BioLegend, 1:100), PE anti-mouse CD103 (clone 2E7, Myltenyi Biotec, 1:20), APC anti-mouse CD172a/SIRPα (clone P84, BioLegend, 1:100), PE anti-mouse CD197/CCR7 (clone 4B12, BioLegend, 1:100), FITC anti-mouse CD207/Langerin (clone caa8-28H10, Myltenyi Biotec, 1:20), PE anti-mouse CD287/TLR7 (clone A94B10, BioLegend, 1:100), APC anti-mouse CD326/EpCAM (clone caa7-9G8, Myltenyi Biotec, 1:100), FITC anti-mouse TCRγδ (clone UC7-1305, BioLegend, 1:1000), PE/Cyanine7 anti-mouse IFN-γ (clone XMG1.2, BioLegend, 1:1000), PE anti-mouse IL-4 (clone 11B11, BioLegend, 1:100), PE anti-mouse IL-17A (TC11-18H10.1, BioLegend, 1:1000).

### CHS model

Mice were sensitized by administration of sensitization reagents to shaved abdominal skin and were challenged with an application of challenge reagents to the ears. The details of the hapten reagents and treatment schedules were as follows. FITC (#F3651, Sigma-Aldrich or F0784, Tokyo Chemical Industry): sensitization on Day 0 and Day 2 with 400 μL/head of 0.5% w/v FITC in acetone/dibutyl phthalate (1:1) and challenge on Day 6 with 20 μL/ear of 0.5% FITC solution; DNFB (#A5512, Tokyo Chemical Industry): sensitization on Day 0 with 25 μL/head of 0.30% (low dose) or 0.74% (high dose) w/v of DNFB in acetone/olive oil (4:1) and challenged on Day 5 with 25 μL/ear of 0.15% (low dose) or 0.44% (high dose) of DNFB solution; OXA (#862207, Sigma Aldrich): sensitized on Day 0 with 150 μL/head of 3% w/v OXA in ethanol and challenged on Day 5 with 25 μL/ear of 1% OXA solution. Ear thickness was measured with a caliper (#005327, Mitutoyo).

### Psoriasis model

A psoriasis model was generated based on a previous study with some modifications ^32^. Briefly, 10 mg of 5% BESELNA cream (#MO 651, Mochida Pharmaceutical), a commercially available imiquimod cream, was topically applied to the ears of the mice once per day for 7 days.

### Flow cytometry

DAPI (#11034-56, Nacalai Tescue) and Fc block (clone 93, BioLegend or clone 2.4G2, Tonbo Biosciences) were added to the cells just before staining with specific antibodies against cell surface molecules. For intracellular cytokine staining, the cells were incubated with 5 μg/mL brefeldin A (#420601, BioLegend), 20 μM monensin (#420701, BioLegend), 50 ng/mL PMA (10008014, Cayman Chemical), and 1 μg/mL ionomycin (#095-05831, Wako) for 12 h, and then treated with the Fixation Buffer (#420801, BioLegend) and Intracellular Staining Perm Wash Buffer (#421002, BioLegend). Fluorescence was detected by a MACS Quant Analyzer (Miltenyi Biotec) or a FACSLyric Analyzer (BD Pharmingen). The data were processed with FlowJo software (Tomy Digital Biology).

### Immunohistochemistry

Ear tissues from CHS model mice, which were fixed in 10% formalin and embedded in paraffin blocks, were subjected to H&E staining.

Epidermal sheets were prepared and analyzed as follows. After 40 min of incubation of the ear skin in 38 mg/mL of ammonium thiocyanate solution at 37 ℃, the epidermal sheet was peeled off from the dermis and was attached to a silane-coated slide glass (#514614, Muto Pure Chemicals). After drying, the glass was incubated in CanGet Signal Solution (#NKB-101, TOYOBO) supplemented with 50 μg/mL of FITC anti-mouse I-A/I-E (clone M5/114.15.2, BioLegend) for 1 h at room temperature in the dark following pretreatment with Ultra-LEAF purified anti-mouse CD16/32 Ab (clone 93, BioLegend) for 30 min. After staining with DAPI and washing with PBS-Tween, the glass was incubated with fluoroshield mounting medium (#AR-6500-01, ImmunoBioScience) and was visualized with an FV3000 microscope (Olympus) at ×20 magnification.

### ELISA

The concentrations of the mouse cytokines, IFN-γ, IL-4, IL-6, IL-12p40, and TNF-α, were determined by ELISA MAX Standard/Deluxe Set-series ELISA kits purchased from BioLegend (#430801 or #430815 for IFN-γ; #431104 for IL-4; #431301 or #431316 for IL-6; #431601 or #431604 for IL-12p40; and #430901 or #430916 for TNF-α).

### Quantification of mRNAs

Total RNA was extracted from cells using the ReliaPrep RNA Cell Miniprep System (#Z6012, Promega) and from skin using RNAzol RT Reagent (#RN190, Cosmo Bio) or ISOGEN (#311-02501, Nippongene). RevaTraAce qPCR RT Master Mix (#FSQ-201, TOYOBO) was used to synthesize cDNA from total RNA by RT. Quantitative PCR was performed on a StepOne Real-Time PCR System (Applied Biosystems) with THUNDERBIRD probe qPCR Mix (#QPS-101, TOYOBO) or THUNDERBIRD SYBR qPCR Mix (#QPS-201, TOYOBO) using the primers listed in Supplementary Table 1.

### Western blotting

The cell lysates were resolved via 10% SDS-PAGE and subsequently transferred to a PVDF membrane. After treatment with primary antibodies, including anti-PU.1 (clone D-19; Santa Cruz Biotechnology), anti-IRF4 (clone M-17; Santa Cruz Biotechnology), anti-IRF8 (clone C-19; Santa Cruz Biotechnology), and anti-β-actin (clone AC-15; Sigma-Aldrich), appropriate peroxidase-conjugated antibodies, such as peroxidase-conjugated anti-mouse IgG (#NA931; GE Healthcare) or peroxidase-conjugated anti-rabbit IgG (#NA934; GE Healthcare), were added as the secondary antibodies to the buffer in which the membrane was soaked. The HRP substrate signals were detected with a LAS-4000 (FujiFilm).

### Plasmid construction

For transient expression experiments, pIRES2-AcGFP1 (Clontech Laboratories) and pIRES2-AcGFP1-Myc-IRF4^15^ were used. In addition, we generated two plasmids for the retrovirus vector, pMXs-IRES-deRed-mIRF4 and pMXs-IRES-dsRed, in the present study. To generate pMXs-IRES-dsRed-mIRF4, we first inserted the *Sal*I site at just downstream of the termination codon of the dsRed cDNA of pIRES2-dsRed-IRF4^18^ using a QuickChange II Site-Directed Mutagenesis Kit (Agilent Technologies), and isolated the *Bgl*II-*Sal*I fragment containing the mIRF4-IRES-dsRed DNA region from the mutated plasmid. By inserting this *Bgl*II-*Sal*I fragment into *Bam*HI/*Sal*I-digested pMXs-IG^33^, we obtained pMXs-IRES-dsRed-mIRF4. We also generated pMXs-IRES-deRed, which is a mock vector of pMXs-IRES-deRed-mIRF4, by removing the mIRF4 cDNA fragment from pMXs-IRES-dsRed-mIRF4 via *Xho*I/*Bam*HI digestion, and subsequent self-ligation of the mIRF4 cDNA-deleted fragment after removing the cohesive ends.

### Electroporation of transient expression plasmids or siRNAs

Ten micrograms of expression plasmid or 20 pmol of siRNA were introduced into 1 x 10^7^ BMDC cells by electroporation using a Nucleofector 2b (Lonza) with an Amaxa Mouse Dendritic Cell Nucleofection Kit (Lonza). The abovementioned pIRES2-AcGFP1-series plasmids were used for transient expression, and the following siRNAs were purchased from Invitrogen and used for knockdown experiments: *Nr4a3* siRNA (MSS207089), *Irf4* siRNA (MSS205499), *Spi1* siRNA (MSS247676), and their appropriate GC content-matched controls from the Stealth RNAi siRNA Negative Control Kit (#12935100).

### Retroviral transfection

Retrovirus vectors were prepared by introducing pMXs-IRES-dsRed, or pMXs-IRES-dsRed-mIRF4 into Plat-E^34^ with Fugene6 Transfection Reagent (#E2691, Promega) as described in our previous studies^35,36^. At 4 and 5 days after transfection, culture supernatants of Plat-E cells were harvested to obtain virus solutions. After developing into DCs by cultivation in GM-CSF-supplemented conditions for 3 days, the BM-derived cells were suspended in a viral vector solution that included 10 μg/mL polybrene (#H9268, Sigma-Aldrich). The tubes containing BM-derived cells in viral vector solutions were kept on ice for 10 min and then centrifuged at 1500 × *g* for 90 min at room temperature to perform spin infection. After infection, the cells were maintained in GM-CSF-supplemented media for an additional 6 days.

### Statistical analysis

A two-tailed Student’s t test was used to compare two samples. To compare more than three samples, one-way ANOVA followed by Tukey’s multiple comparison test or Dunnett’s multiple comparison test was used. *p* values <0.05 were considered significant.

## Acknowledgments

We thank members of the Laboratory of Molecular Biology and Immunology, Department of Biological Science and Technology, Tokyo University of Science, for constructive discussions and technical support.

This work was supported in part by a Grants-in-Aid for Scientific Research (B) 23H02167 (CN) and 20H02939 (CN), a Research Fellowship for Young Scientists DC2 (KN), and a Grant-in-Aid for JSPS Fellows 21J12113 (KN) from JSPS; a Scholarship for a Doctoral Student in Immunology (from Japanese Society for Immunology to NI); a Tokyo University of Science Grant for President’s Research Promotion (CN); the Tojuro Iijima Foundation for Food Science and Technology (CN); a Research Grant from the Mishima Kaiun Memorial Foundation (CN); and a Research Grant from the Takeda Science Foundation (CN).

We greatly appreciate the consideration from Dr. Kimihiko Yasuda, Dr. Masako Yasuda, and the late Ms. Yayoi Yasuda.

## Author contributions

M.Katagiri, N.I., K.N., and S.N. performed the experiments, analyzed the data, and prepared the figures; N.M., N.Y., M.T., M.Kurihara, M.N., M.A., and T.Y.. performed the experiments; T.I., constructed the plasmids; Y.A. supervised the experiments; and C.N. designed the research, supervised the experiments, interpreted the results, and wrote the manuscript.

## Supplementary information

The online version contains supplementary information containing Supplementary Fig. S1, S2, S3, S4, and Table 1.

