## Supplementary figures and images for "NR4A3 deficiency ameliorates contact hypersensitivity and exacerbates psoriasis by regulating gene expression in dendritic cells"

### Supplemental Fig. S1.tif

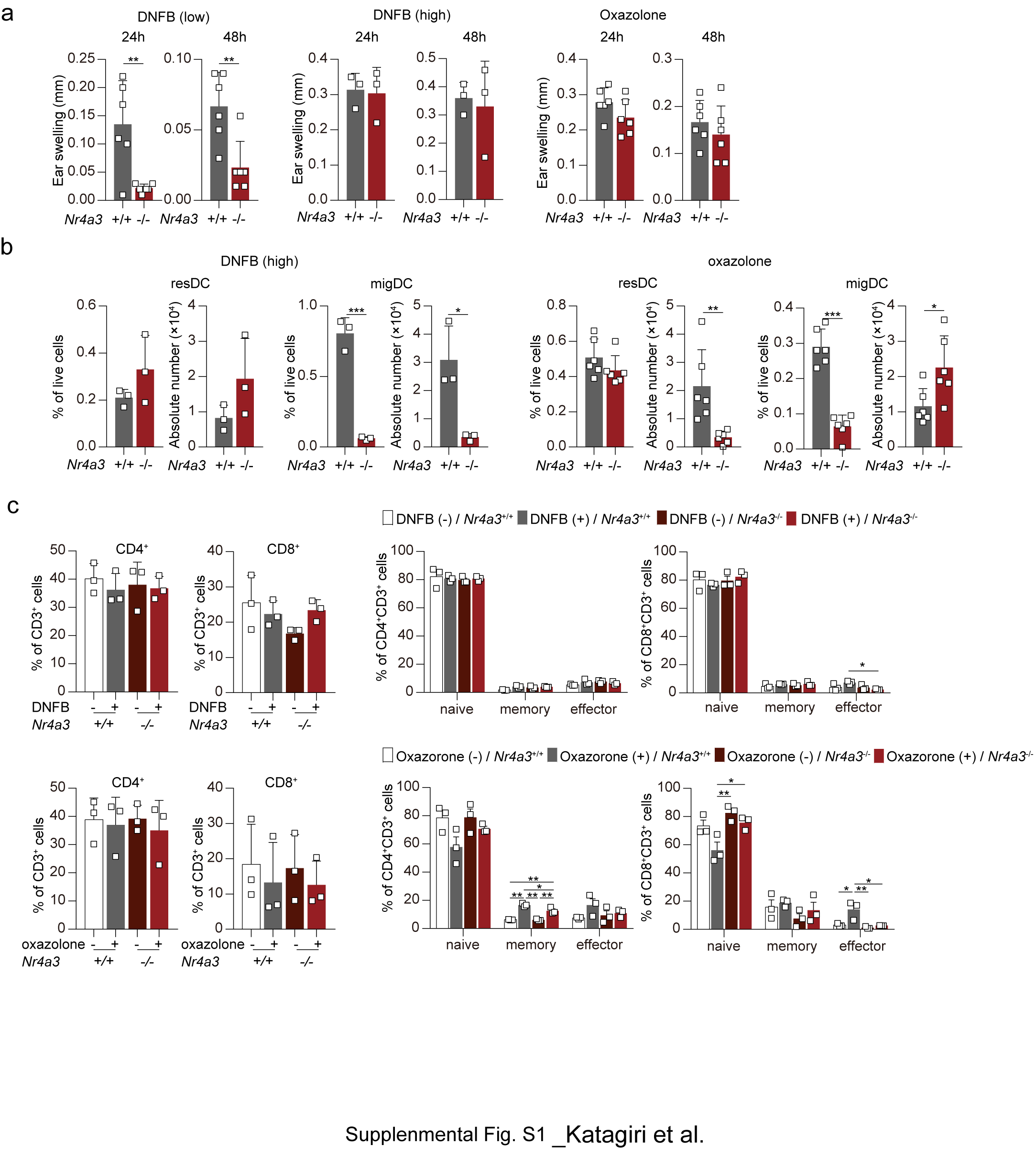

### Supplemental Fig. S2.tif

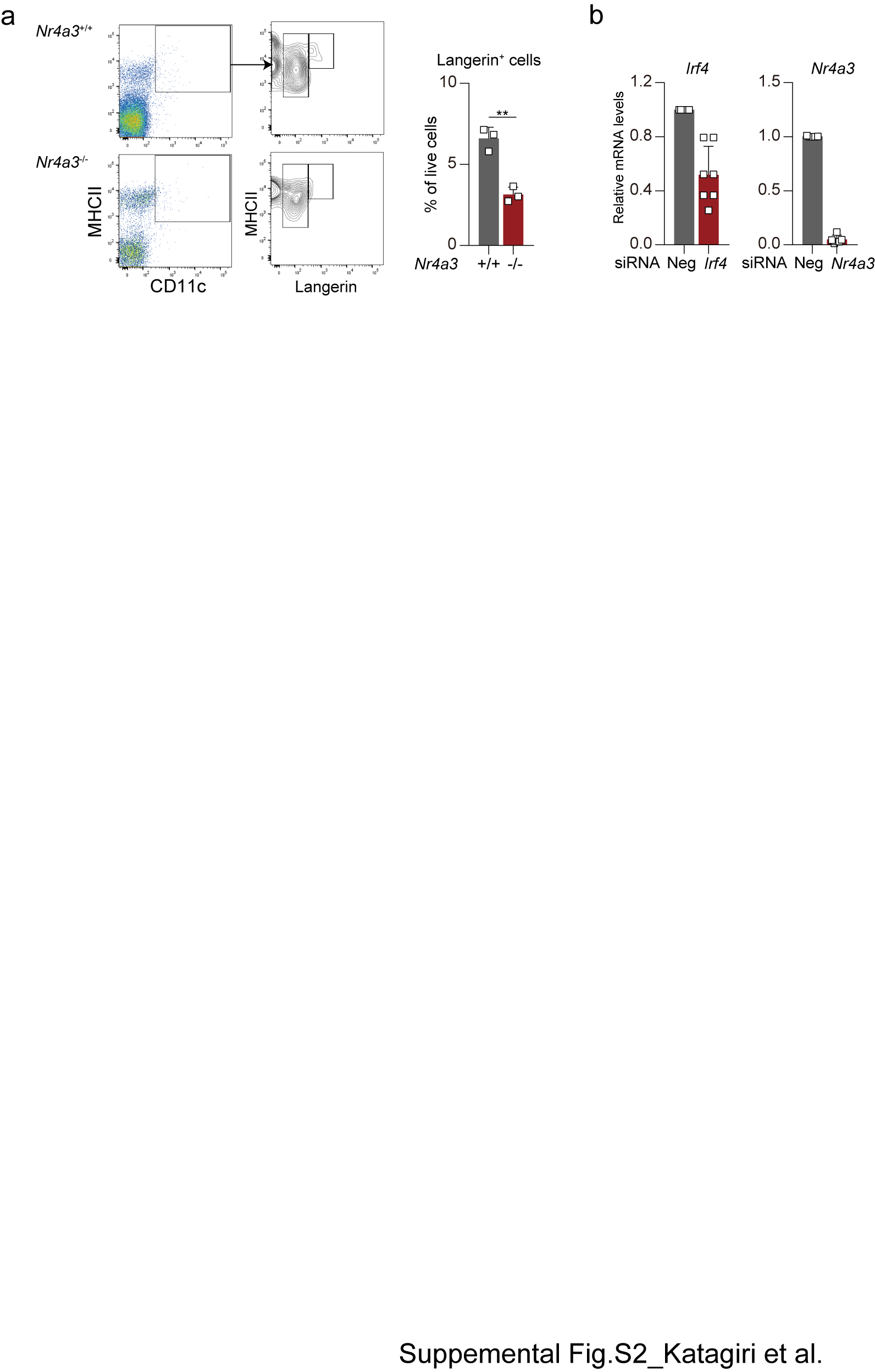

### Supplemental Fig. S3.tif

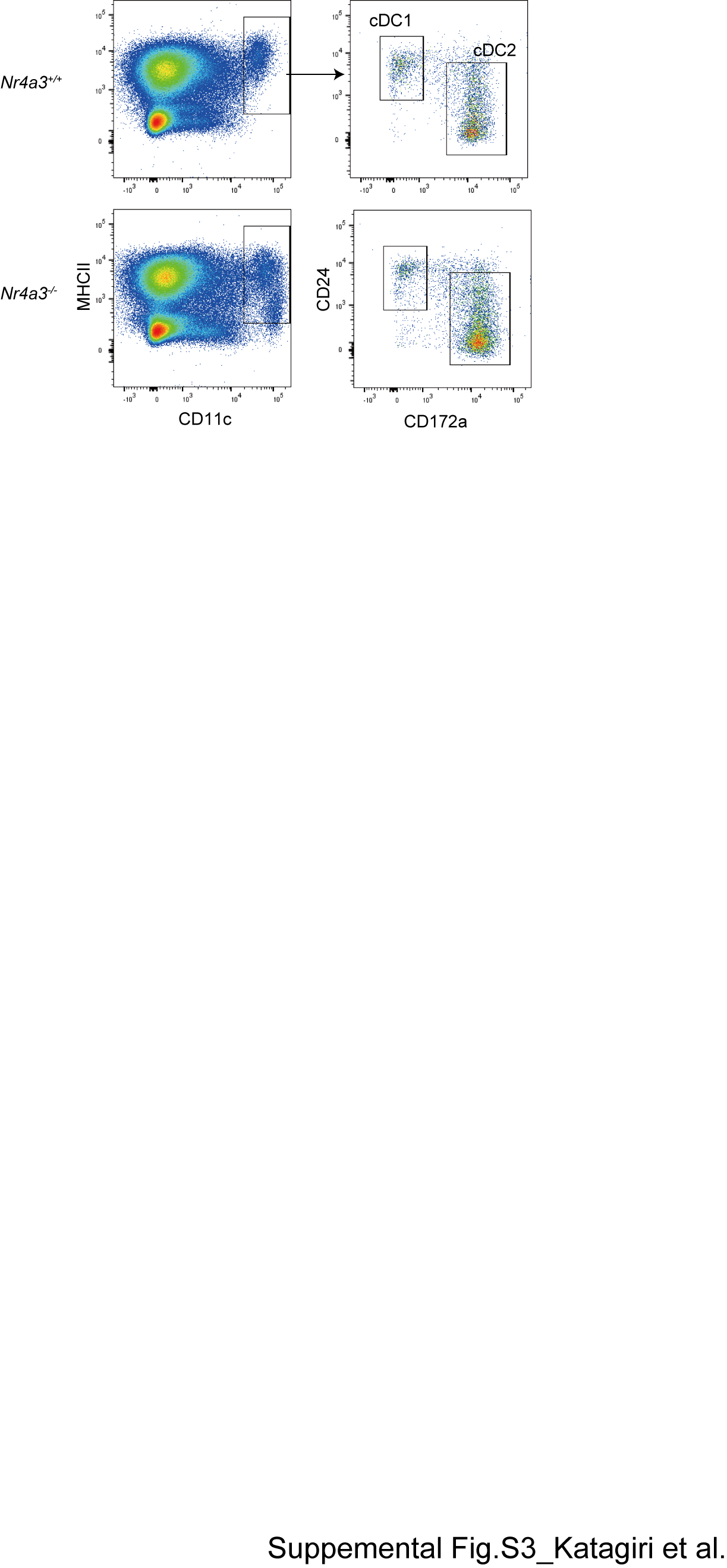

### Supplemental Fig. S4.tif

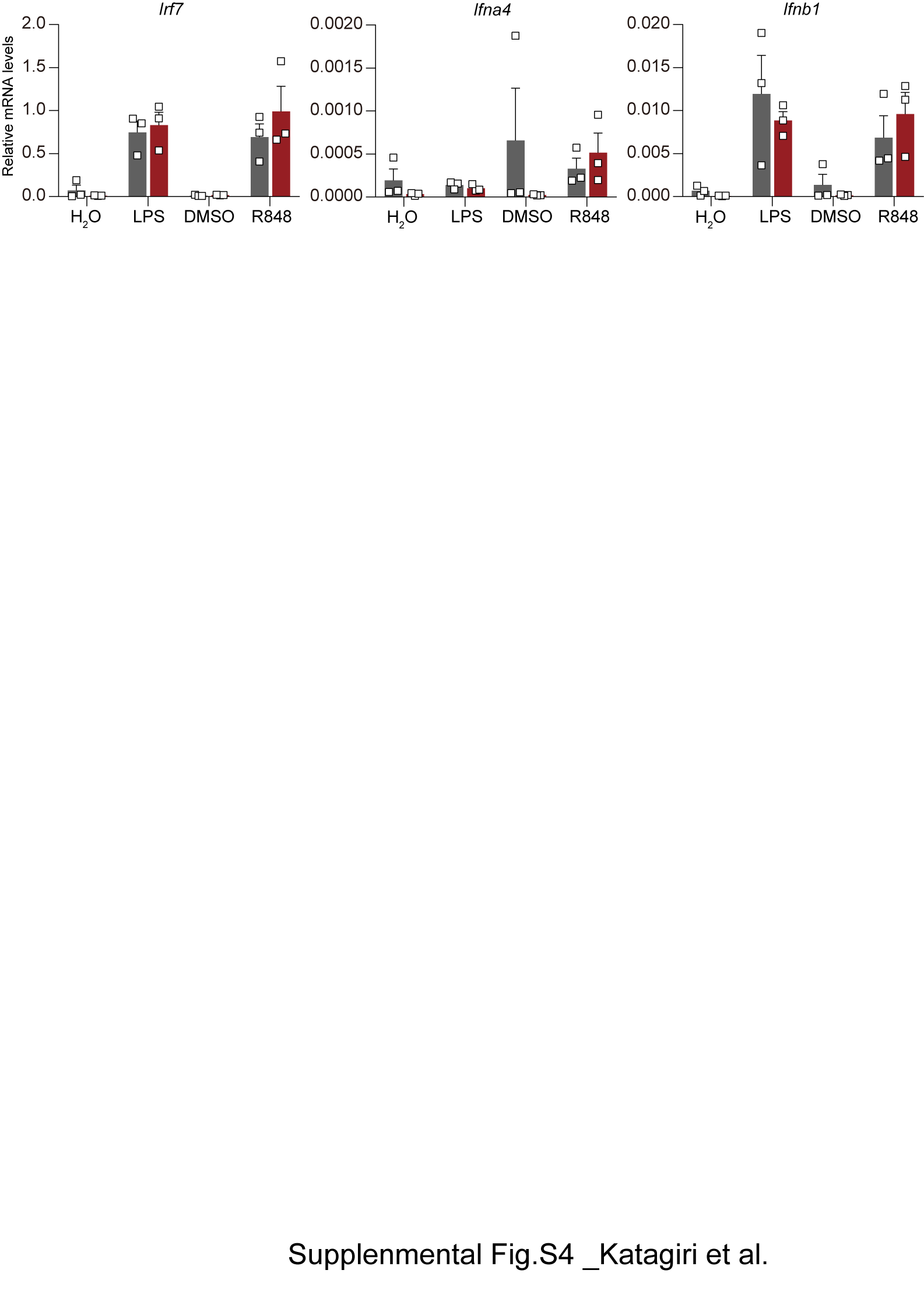
